# Highly efficient genome editing in primary bronchial epithelial cells establishes FOXJ1 as essential for ciliation in human airways

**DOI:** 10.1101/533869

**Authors:** Radu Rapiteanu, Tina Karagyozova, Natalie Zimmermann, Gareth Wayne, Matteo Martufi, Nikolai N Belyaev, Kuljit Singh, Joanna Betts, Soren Beinke, Klio Maratou

**Affiliations:** Target Sciences, GlaxoSmithKline R&D, Stevenage, UK; Refractory Respiratory Inflammation DPU, GlaxoSmithKline R&D, Stevenage, UK

## Abstract

The structure and composition of the bronchial epithelium is altered in respiratory diseases such as COPD and asthma, in which goblet cell hyperplasia and reduced numbers of ciliated cells impair mucociliary clearance. We describe a robust genome editing pipeline to interrogate modulators of primary human bronchial epithelial cell (HBEC) differentiation and function. By employing plasmid- and virus-free delivery of CRISPR/Cas9 to human airway basal cells we achieve highly efficient gene inactivation without the need for positive selection. Genome edited cells are differentiated at air liquid interface (ALI) into a pseudo-stratified epithelium. We focus on profiling ciliation using imaging cytometry coupled to confocal microscopy and immunohistochemistry. To our knowledge, this is the first study to describe highly efficient genome editing of ALI cultured primary HBECs. As proof of concept, we establish that inactivation of the gene encoding the transcription factor FOXJ1 in primary human airway basal cells precludes ciliation in ALI differentiated bronchial epithelia.

## Introduction

The healthy human airway is lined by a pseudo-stratified epithelium containing basal, ciliated and secretory cells, among multiple other cellular subsets (1, 2). A common feature of several chronic airway diseases is the dysregulated induction of secretory cells (which results in excessive mucus production) and the reduction in ciliated cell numbers (which hinders mucus clearance) (3, 4). This leads to obstructed airflow and impaired clearance of pathogens which trigger exacerbations (5). Current standard of care focuses on alleviating disease symptoms such as airway obstruction and inflammation using bronchodilators or steroids, respectively. Whilst these approaches are somewhat effective, they do not address the underlying pathogenic processes. Emerging therapeutic strategies aim to reduce mucus load by altering the properties of mucus (such as viscosity) (6, 7) or limit the production of mucus inducing cytokines (8–10). An alternative way forward would be to target the dysregulated cellular pathways responsible for impaired ciliated cell differentiation. Surprisingly though, given the pervasiveness of airway remodelling in disease, the mechanisms that regulate the homeostasis of the bronchial epithelium are rather poorly understood.

The primary cell based air liquid interface (ALI) system is an established *in vitro* model of human airways that allows the study of differentiation and function of bronchial epithelia (1, 2, 11, 12). Following a 28 day differentiation at ALI, primary human BECs form a pseudo-stratified epithelium containing ciliated and secretory cells (1, 2, 13–15). To facilitate functional studies at ALI, rodent lung samples are predominantly used to source primary cells due to the availability of genetically engineered animals. However, species specific differences pose concerns when studying complex mechanisms. Furthermore, ethical, regulatory, cost and time considerations are making animal studies increasingly difficult. Thus, we sought to develop a genome editing pipeline that would allow us to dissect the molecular mechanisms underlying primary human BEC differentiation and function at ALI.

The ability to edit the genomes of disease relevant, primary human cells has paved new avenues for target validation studies and opened new clinical opportunities. However, efficient genome editing of primary HBECs cultured at ALI has not been previously achieved, due to the fastidious nature of primary HBECs and the intricacies of the ALI differentiation system. CRISPR-Cas9 provides a robust and cost-effective means to target specific genes in mammalian cells. There are multiple ways of delivering Cas9 including plasmid DNA or mRNA transfection or lentiviral vectors. Whilst these methods may very well work in cell lines, they proved suboptimal for our purposes. Firstly, clonal selection and expansion is not possible in primary cells due to their limited proliferative capabilities. Therefore, it is essential to achieve high genome editing efficiencies in bulk populations. Secondly, added complexity arises from maintaining cell health and viability post Cas9 delivery to assure that the delicate downstream differentiation process is not hindered. Furthermore, due to the lengthy ALI differentiation protocol, we reasoned that lentiviral integration is less favourable, as stable Cas9 expression could destabilise HBECs and impact differentiation.

In this manuscript, we describe a fast, robust, cloning-free method to genome edit primary HBECs with high efficiency, without the need for downstream selection of knock-out (KO) cells. Edited HBECs are then differentiated at ALI and ciliation is profiled by imaging cytometry. Thus, we describe a complete pipeline to identify modulators of primary HBEC ciliogenesis and demonstrate that this method can be used to interrogate candidate gene targets.

## Results

### Highly efficient gene disruption in HBECs

The transfection of Cas9 protein pre-complexed with guide RNA (Cas9 <u>r</u>ibo<u>n</u>ucleic <u>p</u>rotein, Cas9 RNP) has been previously shown to offer superior editing efficiencies and reduced toxicity in primary cells (16–21), as compared to plasmid transfection or viral transduction. To test whether Cas9 RNP transfection could be used to inactivate genes in primary human bronchial epithelial cells (HBECs) we targeted the major histocompatibility complex class I (MHC I). MHC I is well expressed on the surface of most nucleated cells and there are excellent flow-cytometric detection reagents available. Therefore, this system could be used to optimise editing efficiencies in most cell types, a procedure we also employed previously (21). Due to the polymorphic nature of the α chain of MHC molecules, we targeted the invariable light chain of MHC I molecules, beta-2-microglobulin (B2M). Depletion of B2M impedes MHC I assembly and targets it for ER associated degradation (ERAD) (22, 23). Therefore, the efficiency of CRISPR/Cas9-mediated depletion of B2M could be quickly and robustly assessed by flow-cytometric analysis of MHC I levels on the surface of HBECs.

Cas9 protein was complexed with tracrRNA plus crRNA targeting the first exon of the B2M gene, and lipofected into primary airway basal cells (Figure 1A). We tested multiple conditions (Figure S1A) and found that complexing 6pmols of Cas9 RNP with 0.75μl lipofectamine and using this mixture to reverse-transfect 50.000 HBECs resulted in the best compromise between MHC I protein depletion and viability (Figure S1A, B). Maintaining viability and optimal growth conditions is a prerequisite for successful HBEC differentiation at ALI. Moreover, we found that extensive handling, temperature variation and manipulation hindered HBEC transfection. By minimising handling times during the transfection protocol, we subsequently achieved B2M gene knock-out in 91% of basal cells (Figure 1B top and middle panel). A clearer separation between knock-out and non-edited cells was observed at 96h post-transfection (Figure 1B, middle panel), as compared to 48h (Figure 1B, top panel). This is due to MHC I protein turnover. B2M depleted cells were cultured at ALI and MHC I depletion was assessed post differentiation (at 36 days). We found that 92% of ALI cultured HBECs did not express MHC I, confirming that the B2M gene modifications were maintained throughout the 28 day differentiation (Figure 1B, lower panel).

**Figure 1.**
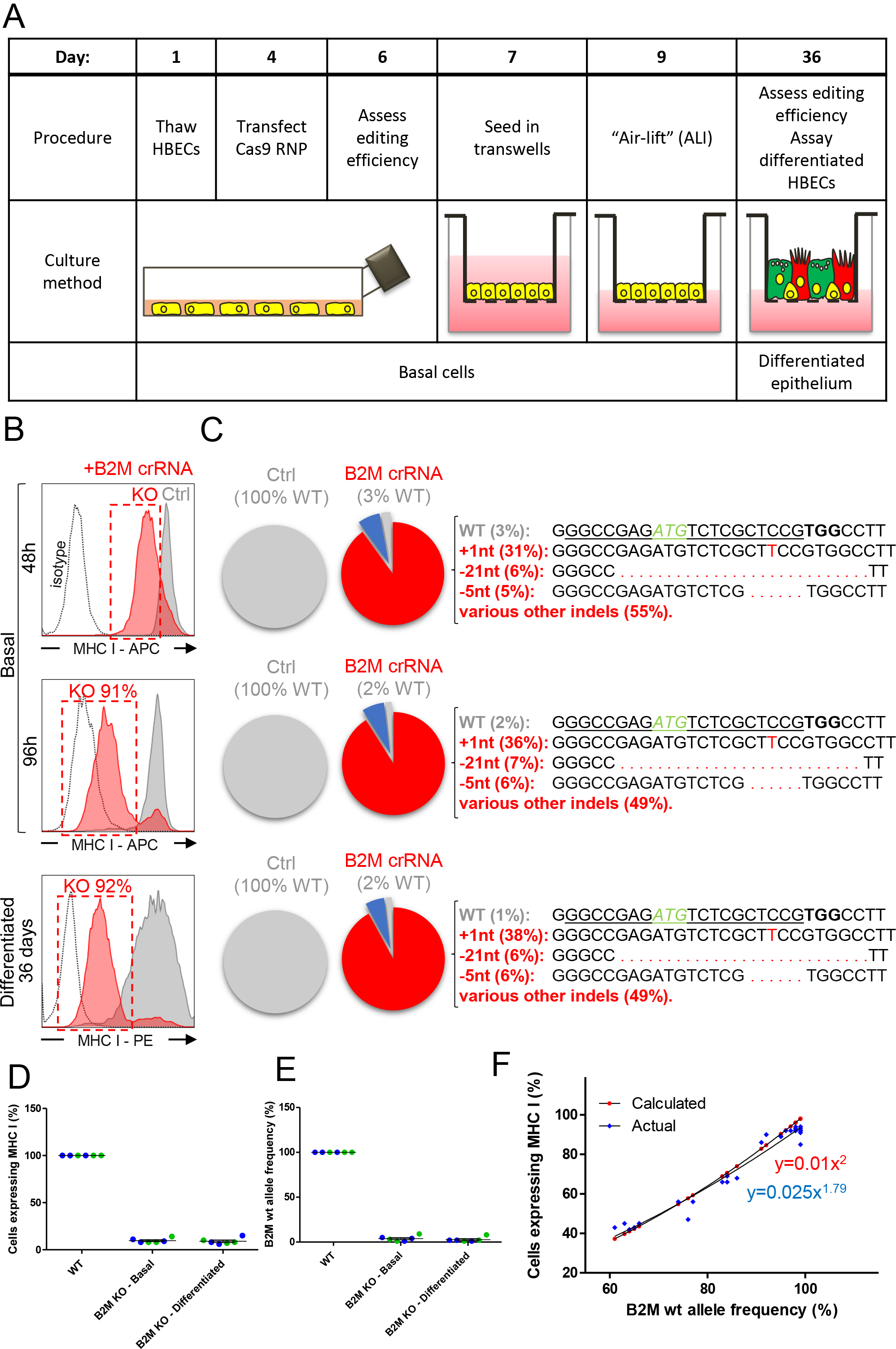
High genome editing efficiencies are maintained throughout HBEC differentiation at ALI. (A) Overview of primary HBEC genome editing and ALI differentiation protocol. (B) Cytofluorometric analysis of MHC I expression in B2M depleted HBECs (red histogram) derived from donor NS838 at multiple time points: 48h, 96h and 35 days post transfection. Ctrl (grey histogram) – non-transfected cells; KO (red dotted gate) – B2M knock-out cell population. (C) MiSeq genotyping of B2M depleted HBECs derived from donor NS838 at 48h, 96h and 35 days post transfection. Pie charts represent the total number of sequenced alleles. Wild-type alleles are marked in grey whilst indels are coloured in red and blue. Top 3 gene alterations (by percentage of total alleles) are shown in red on the right along with their sequence. B2M targeting crRNA sequence is underlined while the PAM is in bolded text. The B2M translation initiation codon is shown in green, italic letters. Cas9 induced sequence modifications are shown in red. (D) Percentage of cells expressing MHC I after B2M gene inactivation. Mean ± SEM from experiments in 2 healthy donors, across 3 technical repeats. Donor NS838 – green data points; Donor NS955 – blue data points. (E) Frequency of wildtype B2M alleles after B2M gene editing. Showing populations described in (D). (F) 2-dimensional plot of wildtype B2M allele frequency versus percentage of cells expressing MHC I, after B2M gene inactivations of varying efficiencies. Red data points and equation represent the theoretical correlation. Experimentally determined correlation shown by blue data points and equation.

To accurately determine the resulting genetic modifications, we used a previously described MiSeq genotyping protocol (21). Briefly, we performed high-throughput sequencing and analysed the data using the freely available online tool OutKnocker (24, 25). Outknocker identifies sequencing reads that map to the referenced, CRISPR-targeted locus. It then outputs the frequencies of wildtype and edited alleles. We illustrate this information via pie charts which represent the total numbers of alleles in the sequenced sample. Wild-type alleles were marked in grey. Edited alleles containing indels (<u>in</u>sertions or <u>del</u>etions) that cause out-of-frame mutations were coloured in red whilst those containing in-frame mutations were coloured in blue. As a control, a wild-type, non-edited population of cells (from the same donor) was sequenced in parallel to assure that no genetic variation occured at that particular locus. MiSeq genotyping of HBECs targeted with B2M crRNA revealed that 97% of alleles were edited 48h post transfection (Figure 1C). When assessing the top 3 sequence modifications we found that 31% of the total number of alleles in the edited population contained a 1 nucleotide insertion, followed by a 21nt deletion and a 5nt deletion that together made up for 11% of total alleles. The editing efficiencies and the distribution of edited alleles was conserved throughout the 28 day differentiation suggesting that there was no selective pressure on specific alleles (Figure 1C).

To test the robustness of the editing protocol, we edited the B2M gene in primary HBECs originating from both healthy (NS838 and NS955) and diseased (C083) donors. We routinely achieved MHC I depletion in 90% of basal cells (Figure 1D and S1C), as assayed by flow-cytometric analysis, and 96% gene editing efficiency (Figure 1E and S1D), as assayed by MiSeq genotyping of the B2M locus. Furthermore, these protein depletion and editing efficiencies were maintained throughout the 28 day differentiation (Figure 1D, E and S1 C, D). The flow cytometry data taken together with the genomic analysis demonstrate that B2M was efficiently edited in primary HBECs derived from multiple donors across multiple technical repeats.

In line with recent studies in other primary cells (16–21), we have also developed an electroporation-based genome editing protocol. We employed this protocol to edit the B2M gene in primary basal HBECs originating from two healthy (NS838 and NS955) donors and routinely achieved 94% gene editing efficiency, as assayed by MiSeq genotyping (Figure S1E). Genome editing efficiencies were maintained throughout ALI differentiation.

We next examined how gene editing efficiencies, assessed by MiSeq genotyping, translated to protein depletion, assessed by flow cytometry. We analysed HBECs edited with various efficiencies and plotted the two metrics in Figure 1F. Given the diploid nature of HBECs, the percentage of cells that lost MHC I expression (i.e. incurred disruptions in both B2M alleles) should equate to the likelihood of having two disrupted alleles per cell, expressed as a percentage. In brief, % KO cells = (% edited alleles)^2^ / 100 (Figure 1F, red series). As expected, our experimental data closely resembled this mathematical equation (Figure 1F, blue series). This correlation becomes useful when targeting proteins toward which robust detection reagents are not readily available, as one can infer the % KO cells from % edited alleles.

### Depletion of FOXJ1 abolishes ciliated cells in primary human ALI cultures

Having determined that efficient genome editing in primary HBECs does not preclude culture at ALI, we next sought to introduce targeted gene disruptions that modulate HBEC ciliation. To assess the proportion of ciliated cells we employed image cytometry, which combines the speed and sensitivity of flow cytometry with the detailed imagery of microscopy. The ImageStream^®^X Mark II (MerckMillipore) produces multiple high-resolution images of every cell directly in flow. This allowed us to accurately determine the number of morphologically distinct ciliated cells in ALI cultures. First, we determined the number of ciliated cells in ALI cultures from non-edited HBECs derived from two healthy donors. Cells exhibiting morphologically distinct cilia (Figure 2A) were selected and plotted on forward scatter (FSC) x side scatter plots (SSC) (Figure 2B). We found that cells derived from donor NS838 routinely produced more ciliated cells in ALI cultures, as compared to HBECs derived from NS955 (Figure 2B, D).

**Figure 2:**
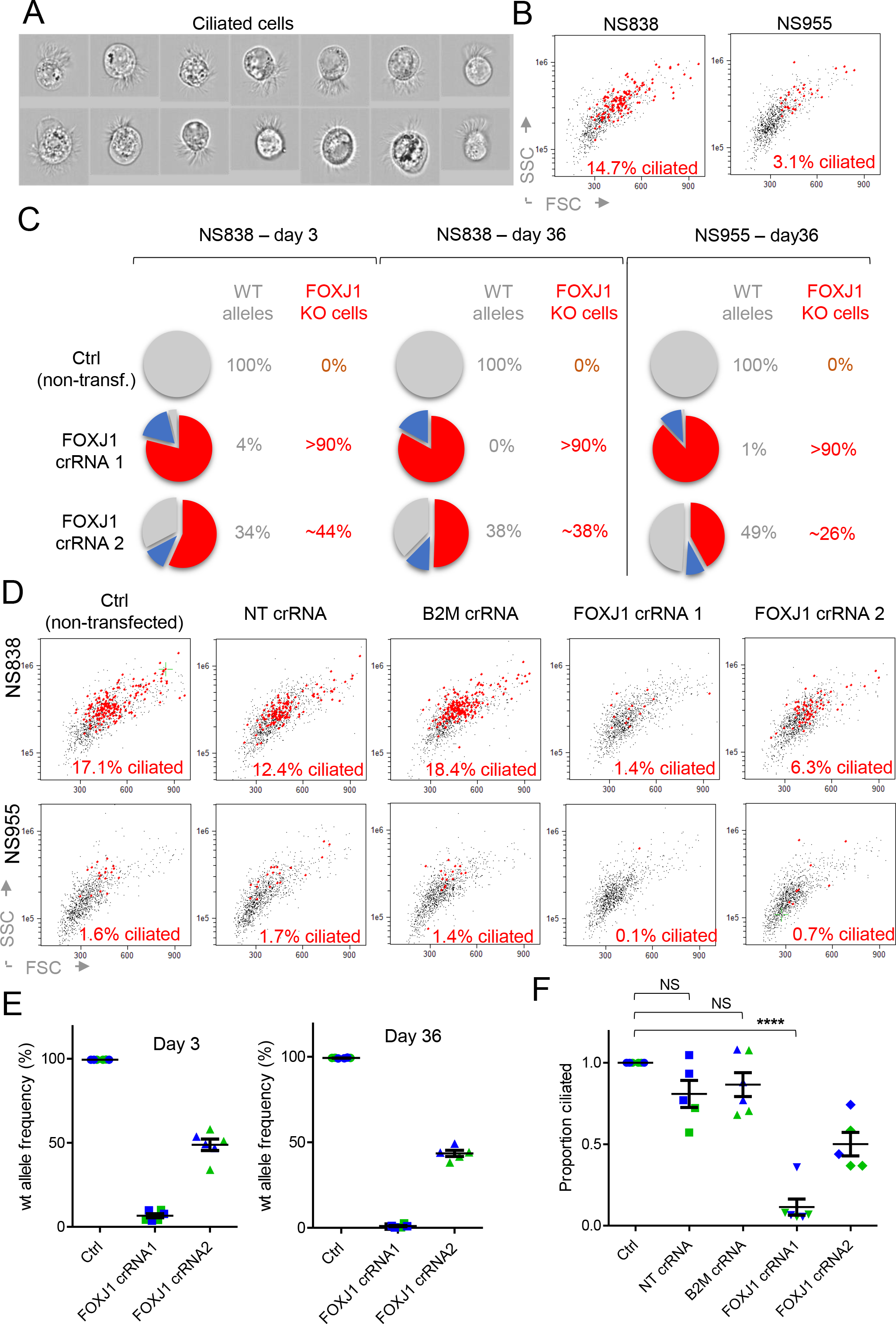
FOXJ1 inactivation impedes HBEC ciliation. (A) Imaging flow cytometric analysis of differentiated HBECs displaying morphologically distinct cilia. (B) Morphologically distinct ciliated cells were plotted on FSC x SSC. ALI cultures were derived from donors NS838 and NS955. (C) MiSeq genotyping of FOXJ1 depleted basal HBECs derived from donor NS838 (day 3 and 36) and NS955 (day 36). Pie charts represent the total number of sequenced alleles. Wild-type alleles are marked in grey whilst indels are coloured in red and blue. %KO was approximated using the previously validated formula: % KO cells = (% edited alleles)^2^ / 100. (D) Imaging flow cytometry analysis of B2M and FOXJ1 depleted HBECs derived from donor NS838 and NS955. Cells displaying morphologically distinct cilia were marked in red. FOXJ1 inactivation efficiencies shown in Figure 2C. Ctrl – non-transfected cells. NT – non-targeting crRNA transfected cells. (E) Frequency of wildtype FOXJ1 alleles after transfection with FOXJ1 crRNA1. Mean ± SEM from experiments in 2 donors across 3 technical repeats. Genome editing efficiencies were assessed at day 3 and 36 post transfection. Donor NS838 – green data points; Donor NS955 – blue data points. (F) Proportion of ciliated cells in non-transfected, NT, B2M KO and FOXJ1 KO HBECs. Mean ± SEM from experiments in 2 donors across 3 technical repeats. The percentage of ciliated cells in control (non-transfected) cultures was normalised to 1 for each technical repeat. NS – non significant, ****p ≤ 0.0001.

As proof of concept, we targeted the transcription factor Forkhead box J1 (FOXJ1), which has been shown to regulate ciliation in animal models [reviewed in (16)]. Two FOXJ1 targeting crRNAs were utilized for this experiment: FOXJ1 crRNA1 which targets the translation initiation site (ATG) and crRNA2 which targets ~35 bases downstream of the ATG (Figure SA, B). Cas9 RNP was transfected into basal HBECs, followed by MiSeq genotyping analysis of editing efficiencies. Three days post transfection, FOXJ1 crRNA 1 generated 96% genome editing efficiency whereas crRNA 2 generated 66% genome editing (Figure 2C and S2B). FOXJ1 KO HBECs, derived from two healthy donors (NS838 and NS955), were then differentiated at ALI. We found that FOXJ1 gene inactivation was maintained throughout the 28 day differentiation suggesting that FOXJ1 depletion is not negatively selected during ALI differentiation (Figure 2C).

We hypothesized that depletion of FOXJ1 would preclude ciliation in ALI differentiated bronchial epithelia. Indeed, efficient disruption of FOXJ1, but not B2M, decreased the proportion of ciliated cells in HBECs derived from two healthy donors (Figure 2D). As expected, the poorer editing efficiencies achieved with FOXJ1 crRNA 2 induced a weaker reduction in ciliated cells (Figure 2D). This observation was consistent in differentiated HBECs derived from both donor NS838 and NS955 (Figure 2D), indicating that the strength of the phenotype was dependent on genome editing efficiency and not due to CRISPR/Cas9 off target effects or transfection or culture artefacts.

To assure the robustness of our findings, we performed B2M and FOXJ1 gene deletions in basal cells derived from NS838 and NS955 (two biological replicates), in three technical replicates. We routinely achieved FOXJ1 editing efficiencies of 93% using crRNA1 and 51% by using crRNA2, as assayed by high-throughput sequencing of the FOXJ1 locus (Figure 2E). We found that FOXJ1 depletion significantly decreased the proportion of ciliated cells in HBECs derived from two healthy donors (Figure 2F). Depending on the nature of our studies we routinely use two differentiation media to culture HBECs at ALI: 3D media and Pneumacult media. Throughout this study, we focussed on 3D differentiation media. However, to assure our findings are valid across two distinct differentiation media we cultured FOXJ1 depleted HBECs (NS838 and NS955) at ALI using Pneumacult media and showed that depletion of FOXJ1 precludes ciliation (Figure S2C).

### FOXJ1 depletion impacts overall organisation, polarization and morphology of human ALI cultures

Having shown that FOXJ1 KO reduces the proportion of ciliated cells by image cytometry, we next explored the functional effects of FOXJ1 depletion using an established orthogonal approach, confocal immunofluorescent microscopy. Apical expression of β-Tubulin IV is a commonly used marker to visualize the distribution of cilia on the apical surface of bronchial epithelia (26–29). Having selected crRNA 1 as the more efficient means of generating FOXJ1 gene inactivation, we generated edits in HBECs derived from donors NS838 and NS955 and differentiated cells at ALI. The resulting bronchial epithelia were fixed, permeabilised and stained for β-Tubulin IV. Due to the fact that an intact differentiated epithelial layer (at day 36) is required for immunofluorescent microscopy, it was only possible to quantify editing efficiencies 3 days post transfection (Figure 3A). However, as described above, FOXJ1 gene deletions are maintained throughout ALI differentiation (Figure 2C, E). We show FOXJ1 depleted epithelia derived from 2 healthy donors exhibit significantly reduced β-Tubulin IV expression (Figure 3B, C, D). Thus, the confocal microscopy data corroborates the image cytometry data to establish that FOXJ1 depletion hinders ciliation.

**Figure 3:**
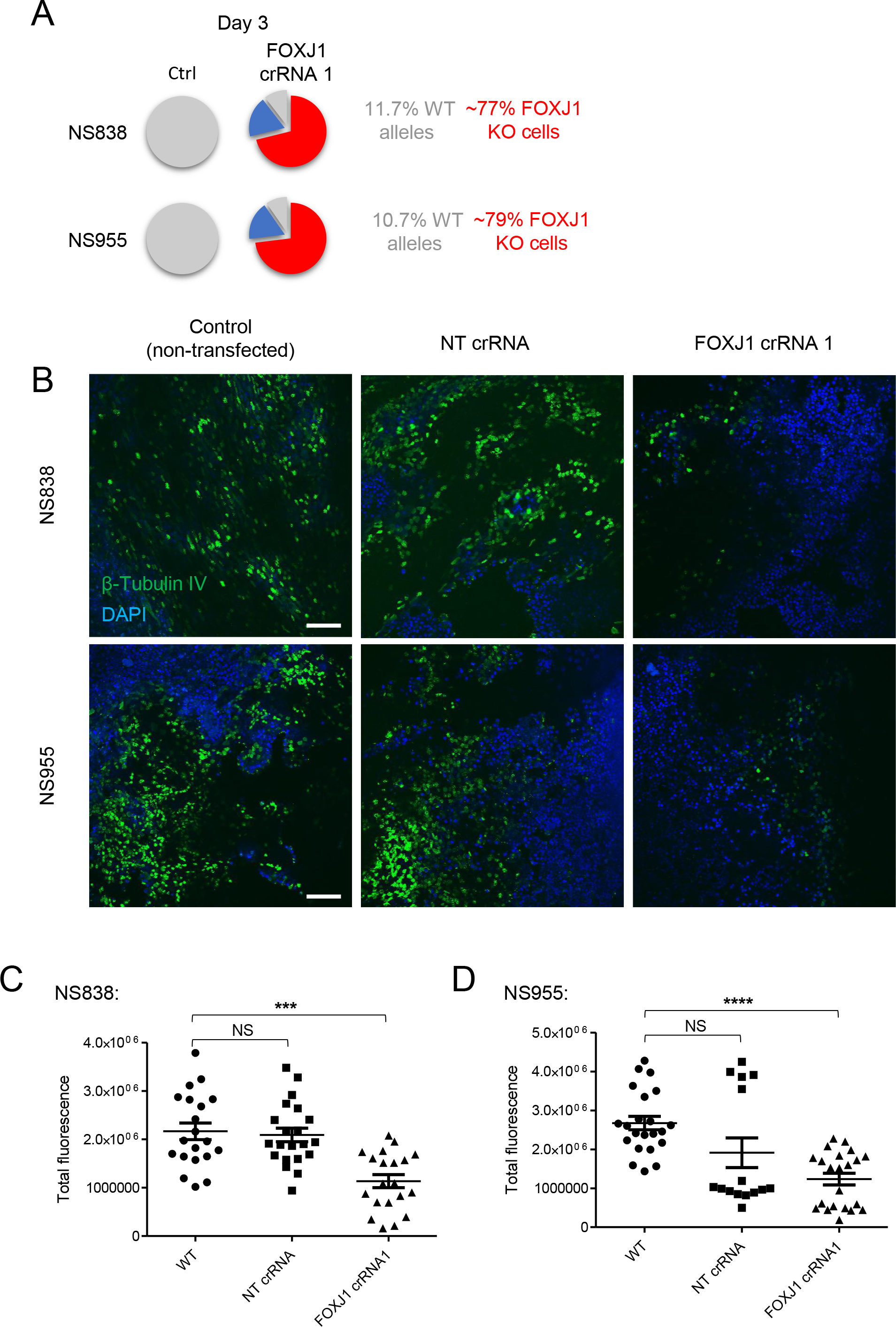
Confocal immunofluorescence microscopy confirms that inactivation of FOXJ1 hinders ciliation. (A) MiSeq genotyping of FOXJ1 depleted basal HBECs derived from donor NS838 and NS955 (day 3). Ctrl – non-transfected cells. %KO was approximated using the previously validated formula: % KO cells = (% edited alleles)^2^ / 100. (B) Confocal immunostaining of β-Tubulin IV in control and FOXJ1 depleted ALI cultures. The brightest β-Tubulin IV field is shown in each case. NT – non-targeting crRNA transfected cells. Scalebar represents 100μm. (C and D) Quantification of total anti-β-Tubulin IV-Alexa 488 fluorescence intensity from confocal images in ALI cultures derived from NS838 (C) and NS955 (D). Mean ± SEM of two independent experiments with 10 fields per condition. NS – non significant, ***p ≤ 0.001, ****p ≤ 0.0001.

To further explore the effects of FOXJ1 depletion on the morphology and ciliation of the pseudostratified bronchial epithelium, we employed histological analysis of sectioned ALI cultures. FOXJ1 KO HBECs, derived from two healthy donors (NS838 and NS955), were cultured at ALI for 28 days prior to fixation and hematoxylin and eosin (H&E) staining. The pseudostratified epithelia originating from non-transfected (control) NS838 cells were polarized and highly organized across 3-5 layers (Figure 4A). Although the cells showed some variation in size and shape, the apical layers maintained a distinctive cuboidal to columnar shape, consistent with previous reports (30). Morphologically distinct cilia were observed on the apical membranes. FOXJ1 depletion in NS838 derived epithelia abolished cilia and, interestingly, induced a squamous morphology (Figure 4A, B). These effects were not observed in cells transfected with NT crRNA, which closely resembled non-transfected cells (Figure 4B). Epithelia originating from non-transfected NS955 cells were organised across 1-3 layers and cells displayed the distinctive cuboidal to columnar shape (Figure 4C). Morphologically distinct cilia were observed on the apical membranes. FOXJ1 depletion in NS955 derived epithelia resulted in a non-ciliated, unpolarised and atrophied epithelium (Figure 4C).

**Figure 4:**
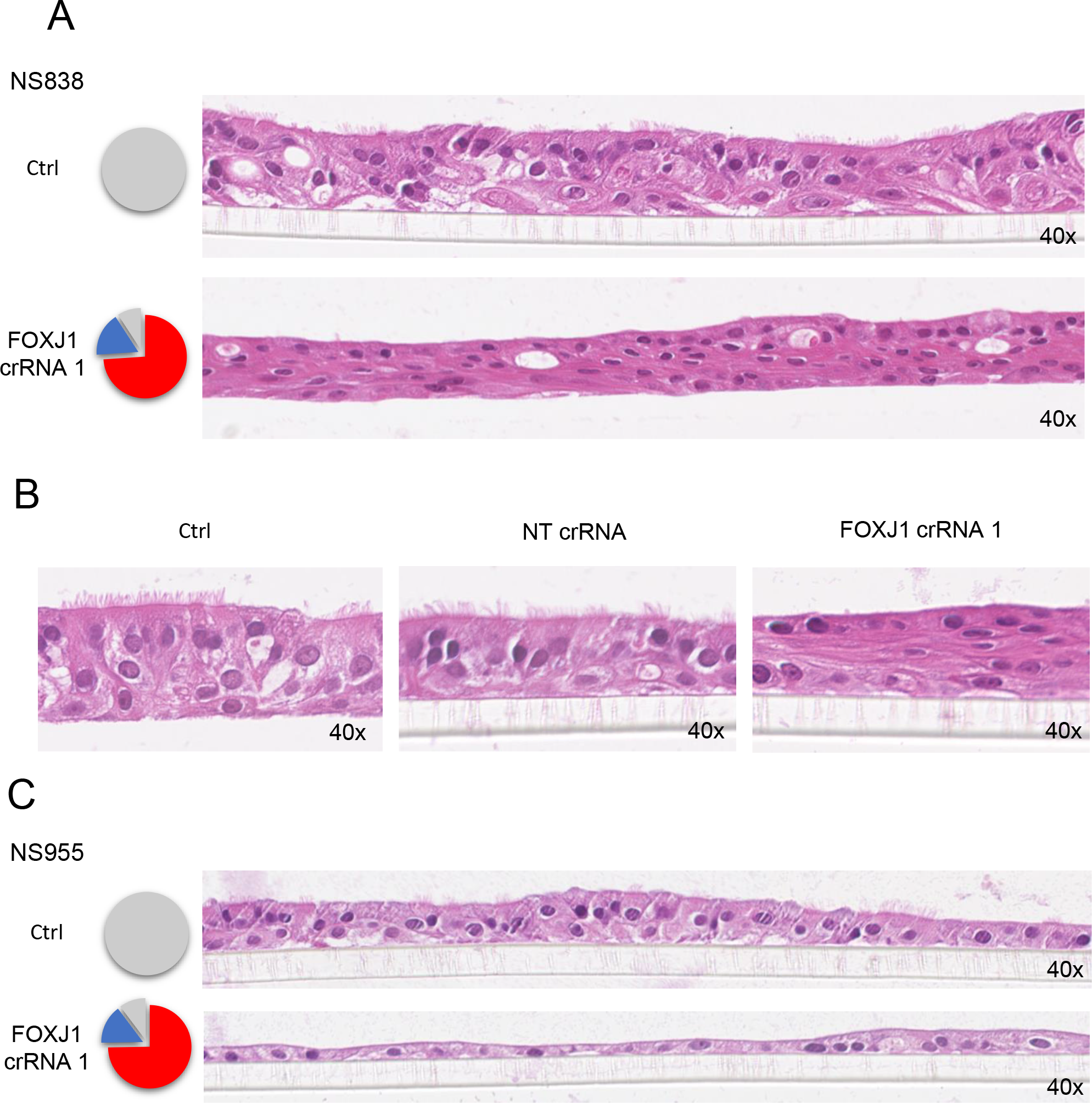
FOXJ1 depletion impacts overall organisation, polarization and morphology of human ALI cultures. (A and B) Histological assessment of ALI cultures derived from NS838. Control and FOXJ1 depleted cells were cultures at ALI for 28 days, fixed and sectioned. Morphology was assessed by H&E staining. Pie charts represent the total number of sequenced alleles at day 3. Wild-type alleles are marked in grey whilst indels are coloured in red and blue. Ctrl – non-transfected cells. NT – non-targeting crRNA transfected cells. All images were acquired at 40x magnification. (C) Histological assessment of ALI cultures derived from NS955.

The IHC and IF data strengthened the imaging cytometry data to establish that disruption of FOXJ1 in primary ALI-differentiated HBECs precludes ciliation. Furthermore, FOXJ1 depletion impacts overall organisation, polarization and morphology of human ALI cultures.

## Discussion

Herein we establish a target validation pipeline that enables the identification of novel factors that modulate primary HBEC differentiation at ALI. The CRISPR/Cas9 RNP-based system allows for highly efficient, cloning- and selection-free, genome editing of primary HBECs. Post ALI differentiation, the effect of specific gene inactivation on ciliation is assayed using image cytometry, a technique which allows the robust assessment of ciliated cell numbers. We demonstrate proof of concept by showing that depletion of FOXJ1 significantly reduces the proportion of ciliated cells in human ALI cultures. To our knowledge, this is the first study to show that FOXJ1 is essential for ciliation in primary human BECs.

FOXJ1 is a well characterised transcription factor required for motile cilia generation in model organisms (31–41). FOXJ1 induces the expression of basal body components; structural, motor and accessory proteins of the axoneme; and vesicular and intraflagellar transport proteins [reviewed in (42)]. Several systematic studies in animal models have been performed to identify the genes regulated by FOXJ1 (36, 38, 40, 41, 43–46). The results were only partially overlapping (43) indicating that the regulatory programme initiated by FOXJ1 varies across species, rendering such studies suboptimal for dissecting human biology. Despite the interest in FOXJ1’s role as a key regulator of motile cilia, research efforts in human cells have been limited to overexpression studies (4, 47) or reporter assays based on FOXJ1 promoter fusions (48). We describe a protocol to KO FOXJ1 in primary HBECs and thus provide a clean background to facilitate further exploratory studies aiming to identify the complete genetic programme of FOXJ1 in human cells. Furthermore, our data show that FOXJ1 depletion induces a squamous morphology of the bronchial epithelium. This suggests that the regulatory programme governed by FOXJ1 may be important not only for ciliation but also for correct pseudo-stratification of bronchial epithelia.

We establish fast and efficient lipofection- and electroporation-based protocols for editing HBECs cultured at ALI. Transient RNP transfection does not require cloning and the gene inactivation is obtained within a week. The high KO efficiencies (over 90%) eliminate the need for KO cell selection, as functional assays can be carried out in pools. Editing of primary human airway basal cells has been previously performed by employing lentiviral transduction (49) or by deriving isogenic cell lines from transfected primary cells (50). However, to our knowledge, this is the first study to describe highly efficient genome editing of primary HBECs (in bulk) which are subsequently successfully differentiated at ALI (without further cellular reprogramming or immortalization).

This pipeline is immediately amenable to functional arrayed CRISPR screens with multi-parametric endpoint analyses: ciliation profiling and epithelium morphology imaging, cytokine production, pathway activation, barrier integrity assays [e.g. transepithelial/transendothelial electrical resistance (TEER)], mucus quantification, transcriptomics, proteomics, etc. Human Rhinovirus or Moraxella catarrhalis challenges could also be integrated in the assay. Edited HBECs could also be co-cultured with immune cells (macrophages, dendritic cells, neutrophils) to better mimic the physiology of the bronchial epithelium. Furthermore, the high editing efficiencies open up the possibility for simultaneously editing multiple genes in primary HBECs to model complex genetic landscapes or study genetic redundancy. In our hands, non-essential gene inactivation is maintained throughout the prolonged ALI differentiation. A potential limitation of this system is the study of essential genes, whose depletion may be selected against during ALI differentiation.

We also show that high genome editing efficiencies were achieved in primary HBECs derived from a COPD donor (Figure S1C, D). This paves the way towards functional studies in primary diseased epithelia (COPD, asthma or cystic fibrosis). Nevertheless, in our experience, ALI differentiation of HBECs derived from diseased donors is rather variable and very much donor-dependent. Therefore, before attempting genome editing, donor quality control experiments should be performed.

In conclusion, we establish an efficient CRISPR/Cas9-based pipeline to identify potential targets that modulate primary human bronchial epithelial cell ciliation at air liquid interface. These methodologies will enable functional studies of the dysregulated cellular pathways responsible for impaired ciliated cell differentiation and mucus clearance in respiratory diseases. Furthermore, we show for the first time that depletion of FOXJ1 in primary human BECs impedes their differentiation into ciliated bronchial epithelia.

## Materials and Methods

### HBECs

Primary HBECs were sourced from Lonza with permission for use in research applications by informed consent. HBECs were obtained from healthy donors NS838 (Cat CC-2540, Lot 0000495838, 32 year, non-smoker), NS955 (Cat CC-2540, Lot 0000410955, 29 year, non-smoker) and NS081 (Cat CC-2540S, Lot 0000436081, 66 year, smoker); and diseased donor C083 (Cat 00195275S, Lot 0000436083, 59 year, smoker, COPD patient). The human biological samples were sourced ethically and their research use was in accord with the terms of the informed consents under an IRB/EC approved protocol.

### HBEC culture and differentiation

Basal HBECs were cultured in antibiotic-free BEGM media (Lonza, CC-3170). Untransfected and genome edited basal cells were trypsinized (Lonza, CC-5034), spun down for 4min at 180xg and 200,000 cells were resuspended in 200μl BEGM and seeded in individual, collagen coated, apical wells of HTS transwell plates (Corning, 3378). 1ml of BEGM media was added to the basolateral well.

After 48h, media was aspirated from the apical well to create the ALI and the transwells were transferred to 24 well plates containing 1ml fresh differentiation media (either 3D or Pneumacult media). 3D media was used throughout this study, expect where clearly specified otherwise. 3D media was made up as a 1:1 mixture of SABM (Lonza, CC-3119) and DMEM (Gibco, 21969-035) to which SAGM supplements (Lonza, CC-4124) were added, omitting retinoic acid. Retinoic acid was added fresh, prior to media changes. Both Pneumacult and 3D media were kept antibiotic free. PneumaCult-ALI Medium was made up as per manufacturer’s recommendations (Stemcell Technologies, 05001).

Cells were cultured at ALI for 28 days, with differentiation media changed every two days. The liquefied mucus produced by cells cultured in 3D media was discarded from apical wells during media changes. In our hands, HBECs differentiated in Pneumacult produce a thick, viscous mucus which could not be removed. Pneumacult cultured cells were washed at day 14 by adding 200μl warm Pneumacult media to the apical well.

### CRISPR/Cas9 delivery (lipofection)

CRISPR/Cas9 RNP reagents were sourced from IDT: AltR S.p. Cas9 Nuclease 3NLS (1074182); tracrRNA (1072534); crRNAs were custom made: B2M [GGCCGAGATGTCTCGCTCCG, (51, 52)], FOXJ1 g1 (GAGAGTCCCCGCAGACATGG), FOXJ1 g2 (CCGGCCCGGCTCCCGAGAGG). Lipofectamine CRISPRMAX Cas9 (CMAX00015) was sourced from ThermoFisher. Transfections were carried out in 24 well plates.

HBEC vials (~800.000 cells/vial) were thawed in a 37LC water bath for 80sec. Cells from each vial were then transferred to individual T75 flasks containing 25ml of warm BEGM media and expanded for 3-4 days at 37°C, 5% CO2, with media changes every 2 days. Upon reaching 70% confluency, basal HBECs were washed in HBSS (Lonza, CC-5034), trypsinized (Lonza, CC-5034) and 50.000 cells were resuspended in 500ul BEGM in wells of 24 well plates. Plates were placed in the incubator while the RNP-lipofectamine complex was being prepared. We found that the sooner the cell suspension was transfected, the better the editing efficiencies.

tracrRNA and crRNA were resuspended in Duplex Buffer (IDT, 11-01-03-01) to 100μM and annealed in Duplex Buffer to a final concentration of 10μM. The mixture was heated to 90°C for 3min and cooled down at RT. 6pmoles of AltR S.p. Cas9 were diluted in PBS to 10uM and assembled with the crRNA-tracrRNA hybrid in a molar ratio of gRNA: Cas9 = 2:1, thus forming the Cas9 RNP complex. The complex was diluted in Opti-MEM I (gipco, 11058-021) to 24μl and 1μl Plus Reagent (Thermo, CMAX00015) was added prior to a 10min incubation. In the mean-time 0.75μl lipofectamine CRISPRMAX (Thermo, CMAX00015) was diluted in Opti-MEM to a final volume of 25μl and incubated 7min at room temperature. The lipofectamine solution was the mixed with the RNP solution and incubated for a further 7min at RT. The complex was then transferred to HBECs and plates were transferred to the incubator. Media was exchanged 16h post transfection with warm, pH equilibrated BEGM.

### CRISPR/Cas9 delivery (electroporation)

tracrRNA and crRNA were resuspended in Duplex Buffer (IDT, 11-01-03-01) to 100μM and annealed to a final concentration of 50μM. The mixture was heated to 90°C for 3min and cooled down at RT. 90pmoles of AltR S.p. Cas9 were assembled with 450pmoles crRNA-tracrRNA hybrid (molar ratio of gRNA: Cas9 = 5:1) resulting in a 10.5μl volume solution to which 0.9μl of electroporation enhancer (100μM; IDT, 1075916) were added. This complex is stable and was left at RT while cells were prepared. HBEC were thawed and briefly expanded as described above. Upon reaching 70% confluency, basal HBECs were washed in HBSS (Lonza, CC-5034), trypsinized (Lonza, CC-5034) and 200.000 cells were resuspended in 8.6μl P3 Primary Cell Nucleofector Solution (Lonza, V4XP-3032). The cell solution was then mixed with the RNP solution, transferred to a well of a 16-well Nucleocuvette Strip (Lonza, V4XP-3032) and electroporated using the CM-113 program on the Amaxa 4D Nucleofector (Lonza). Electroporated cells were then swiftly transferred to 6 well plates containing 2mL pre-warmed BEGM and incubated at 37°C.

### MiSeq genotyping

Primers to amplify the targeted loci were ordered from IDT and targeted the following regions (locus targeting region is underlined): B2M (ACACTCTTTCCCTACACGACGctcttccgatct<u>GGCTTGGAGACAGGTGACGGT</u>, TGACTGGAGTTCAGACGTGTGctcttccgatct<u>AGCACAGCGAGGGCCACAGAGG</u>); FOXJ1 (ACACTCTTTCCCTACACGACGctcttccgatct<u>TGAACCTGGCACCTGGTGGTAG</u>, TGACTGGAGTTCAGACGTGTGctcttccgatct<u>ACATACTTATTCGGAGGAGGCGC</u>). Primers to attach barcodes (NNNNNNNN) and Illumina adaptors were ordered from IDT: AATGATACGGCGACCACCGAGATCTACACNNNNNNNNACACTCTTTCCCTACACGACG CT, CAAGCAGAAGACGGCATACGAGATNNNNNNNNGTGACTGGAGTTCAGACGTGTGCT. The MiSeq genotyping protocol was previously described (21, 24). Briefly, cell pellets were lysed in direct lysis buffer [0.2mg/mL proteinase K, 1mM CaCl_2_, 3mM MgCl_2_, 1mM EDTA, 1% Triton X-100, 10mM Tris (pH 7.5)] and incubated 10 min at 65°C and 15 min at 95°C. The lysates were used as PCR templates to amplify the targeted loci. 10% of the PCR product was used as template for a second run of amplification to attach barcodes and Illumina adaptors. The resulting PCR product was quality controlled on a 4200 TapeStation (Agilent), purified using Agencourt AMPure XP (Beckman Coulter, A63882) and prepared for MiSeq sequencing according to manufacturer’s instructions (Illumina). Indel frequencies were then processed using Outknocker, freely available at http://outknocker.org/ (24). The wildtype and indel (in-frame and out-of-frame) allele frequencies reported by the algorithm were then used to generate pie-charts to visualize editing efficiencies.

### Flow-cytometry

Basal cells were trypsinized (Lonza, CC-5034) and stained with anti-human-MHCI-APC (Biolegend, anti-HLA-ABC, clone W6/32, 311410). Cells differentiated at ALI were detached from transwells using StemPro Accutase Cell Dissociation Reagent (Gibco, A11105-01). Apical wells and basolateral wells were washed twice in PBS and 200μl Accutase were added to the apical well and 500μl to the basolateral well. Cells were incubated at 37□C for 10-15min. Cells were then gently pipetted up-and-down 5 times and incubated for a further 10min. Dissociated cells were then transferred to 1.5ml tubes containing 200μl fresh Accutase. Cells were then pipetted up-and-down gently for 10 times and more vigorously for a further 5 times using a 1000μl pipette tip. Cells were then counted and blocked in Human TruStain FcX (Biolegend, 422301) for 5min at 4□C. Cells were washed and resuspended in 100 μl cold differentiation media with 2μl anti-MHCI (Biolegend, anti-HLA-ABC, clone W6/32, 311406). After a 30min incubation at 4□C, cells were stained with LIVE/DEAD Fixable Yellow Dead Cell Stain Kit (ThermoFisher, L34959) as per manufacturer’s protocol. Stained cells were washed twice in 500ul cold differentiation media and assayed on a 4 laser CytoFLEX S Flow Cytometer (Beckman Coulter, B75408). Data was processed in FlowJo V10: viable cells were gated by FSC-A x SSC-A; Single cells were gated by FSC-A x FSC-W; LIVE/DEAD yellow negative cells were then selected for MHC I expression analysis.

### Image-cytometry

For ciliation profiling, differentiated cells were collected as described above, resuspended in cold differentiation media and assayed on the ImageStreamX Mark II Image Cytometer (Merck, 100220). Data was processed in Ideas 6.2. Cells displaying morphologically distinct cilia were marked in red on forward scatter (FSC) x side scatter (SSC) plots and their frequency was then exported to GraphPad Prism 5 and statistical significance was evaluated using paired, two-tailed t-tests.

### Confocal microscopy

Media was aspirated from the apical surface of the transwells containing differentiated HBECs (day 36) and transwells were washed in PBS for 1 minute. ALI epithelia were fixed by transferring the transwell to 24 well plates containing 600μl Parafix (Pioneer Research Chemicals). 200μl Parafix was added to the apical surface of the transwell and incubated 15 minutes at RT. Fixed cells were then washed twice in PBS and permeabilises with Triton X-100. Post PBS washes, 50 μl β-Tubulin IV (diluted 1:100 in PBS; Sigma, T7941) were added to the apical surface of the transwell and incubated over night at 4□C. Cells were washed and then incubated with donkey α-mouse Alexa 488 (Thermo, A-21202). Post PBS washes, cells were incubated with DAPI (Thermo, D1306) and washed again in PBS. The transwell membrane (containing the fixed, stained HBECs) was cut with a scalpel and mounted onto the microscope slide. Mounting medium (Thermo, P36930) was applied and a cover slip was applied on top. Edges were sealed with varnish and slides were analysed on a Leica TCS SP8 confocal microscope. Total fluorescence was quantified using ImageJ, the data exported to GraphPad Prism 5 and statistical significance was evaluated using paired, two-tailed t-tests.

### Histology

Media was aspirated from the apical surface of the transwells containing differentiated HBECs and cells were fixed in Parafix for 30 min at RT as described above. Transwell membranes were excised and processed overnight through to paraffin wax and subsequently placed into a wax mould. 4μm sections were cut from each paraffin wax block and stained with haematoxylin and eosin (nuclei were stained dark blue, cytoplasm pink). Slides were processed and were digitally scanned using the Nanozoomer Image Analyzer.

