## Supplementary figures and images for "Highly efficient genome editing in primary bronchial epithelial cells establishes FOXJ1 as essential for ciliation in human airways"

### Supplementary figures S1 and S2

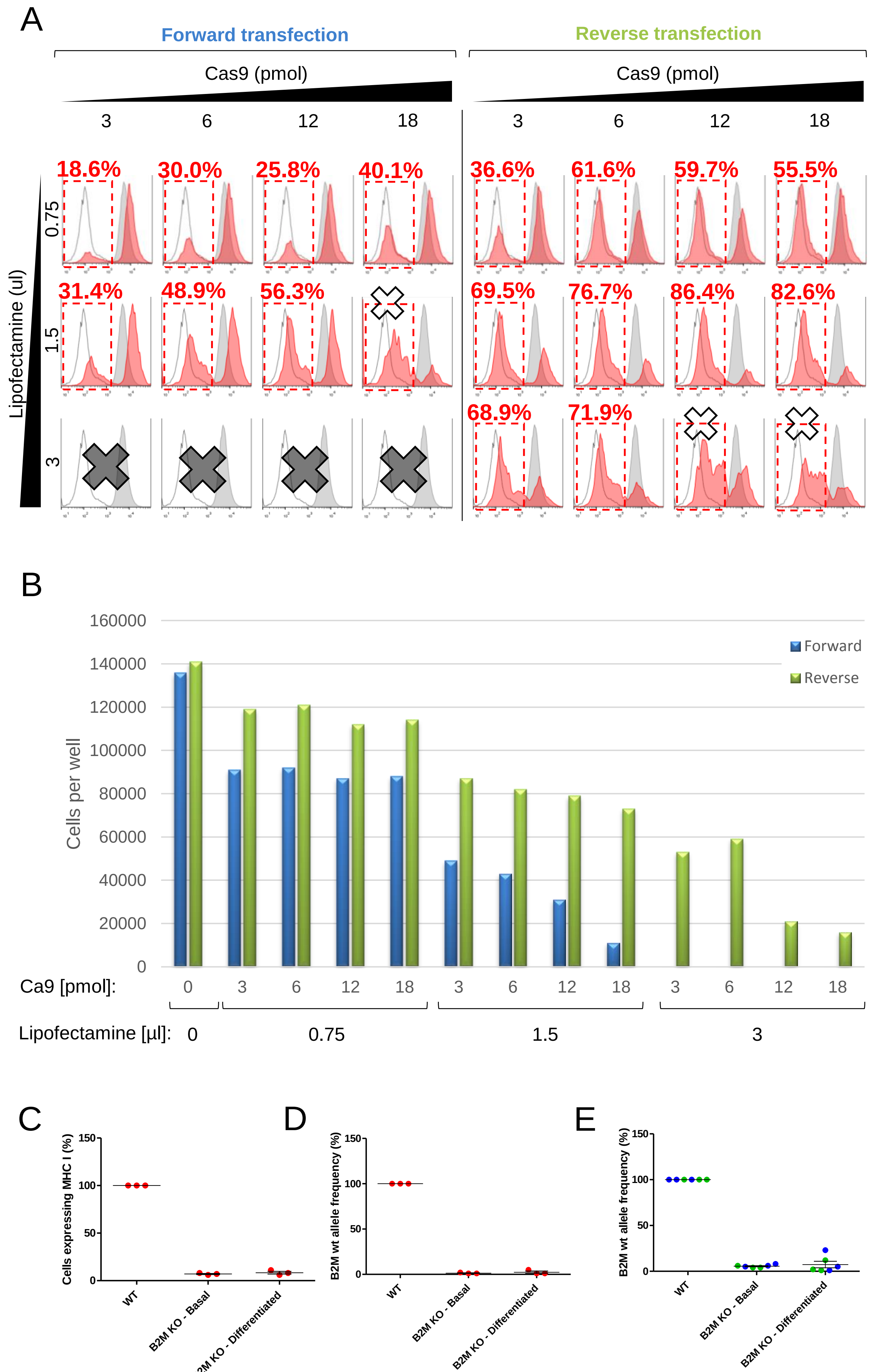

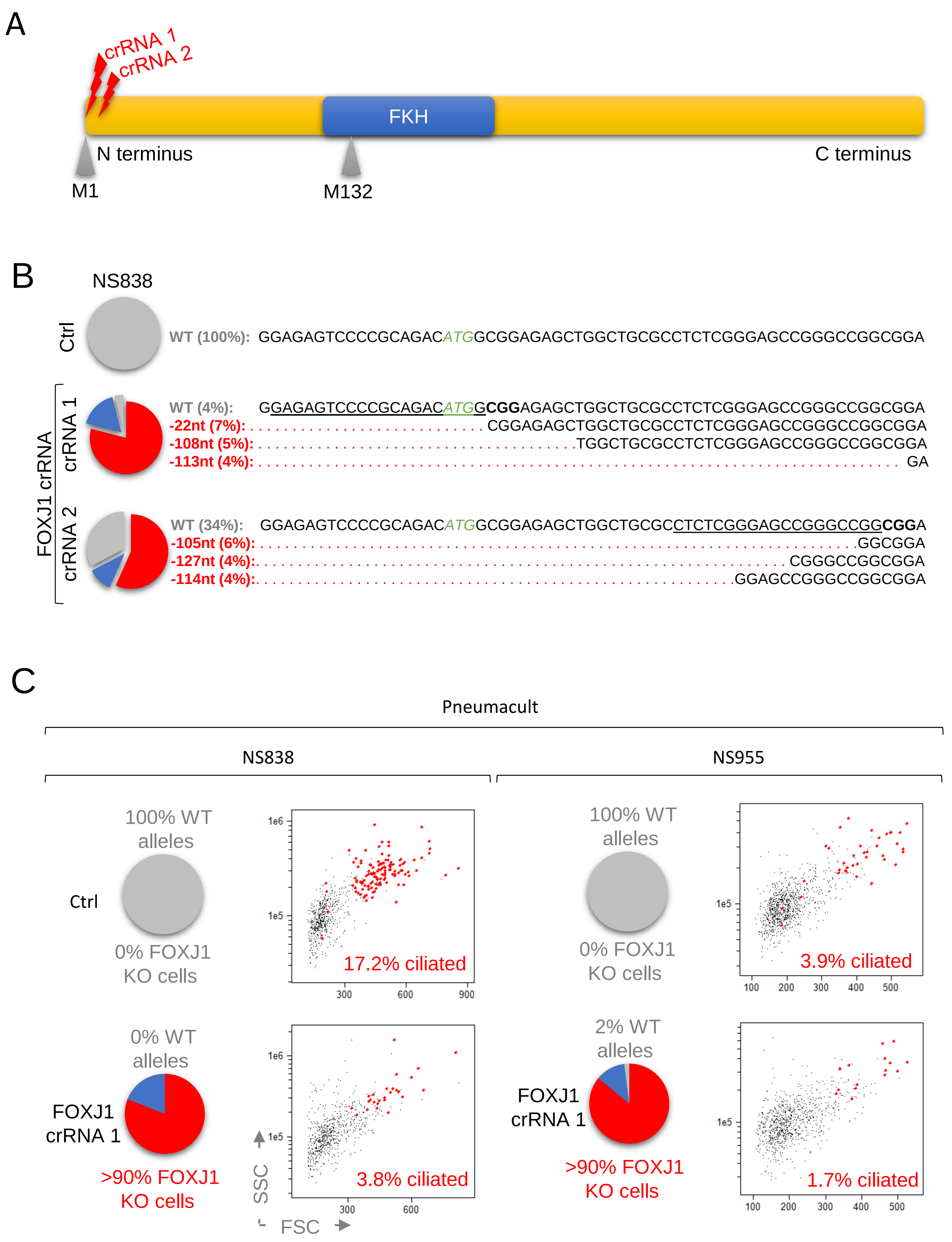
