## Supplementary figure legends for "Highly efficient genome editing in primary bronchial epithelial cells establishes FOXJ1 as essential for ciliation in human airways"

**Figure S1. Optimizing gene editing in HBECs**

(A) Cytofluorometric analysis of MHC I levels in B2M inactivated cells (donor NS081), 72h post transfection. Multiple conditions were tested and the percentage of MHC I depleted cells achieved in each are shown in red. In forward transfections, cells were seeded 24h before transfections and allowed to attach. In reverse transfections, cells were detached and transfected in suspension. Hollow crosses denote samples with fewer than 1000 recorded events. Filled crosses represent samples in which no cells could be recovered.

(B) Survival of HBECs was determined by NucleoCounter® NC-200™ cell counts 72h post transfection.

(C) Percentage of diseased HBECs expressing MHC I after B2M gene inactivation. HBECs were derived from COPD donor C083 and tested across 3 technical repeats.

(D) Frequency of wildtype B2M alleles after B2M gene editing, in HBECs derived from COPD donor C083. Showing 3 technical repeats.

(E) Frequency of wildtype B2M alleles after B2M gene editing using the electroporation protocol. Mean ± SEM from experiments in 2 healthy donors, across 3 technical repeats. Donor NS838 – green data points; Donor NS955 – blue data points.

**Figure S2. MiSeq genotyping of FOXJ1 locus; FOXJ1 KO abolishes ciliation in Pneumacult-differentiated epithelia**

(A) Schematic representation of FOXJ1 and its DNA binding forkhead (FKH) domain. Showing the two crRNAs used to inactivate the gene. Displaying the first (translation initiation site) and second (in frame) methionine residues.

(B) MiSeq genotyping of FOXJ1 depleted basal HBECs derived from donor NS838 (day 3). Pie charts represent the total number of sequenced alleles. Wild-type alleles are marked in grey whilst indels are coloured in red and blue. Top 3 gene alterations (by percentage of total alleles) are shown in red on the right along with their sequence. FOXJ1 targeting crRNA sequences are underlined while the PAMs are in bolded text. The FOXJ1 translation initiation codon is shown in green, italic letters. Cas9 induced sequence modifications are shown in red.

(C) FOXJ1 depletion in ALI cultures differentiated using Stemcell Pneumacult media. Imaging flow cytometry analysis of FOXJ1 depleted HBECs derived from donor NS838 and NS955. Cells displaying morphologically distinct cilia were marked in red. FOXJ1 editing efficiencies are represented on pie charts.
